# An aphid resistant wheat variety reduces the transmission of Barley Yellow Dwarf Virus (BYDV) by *Rhopalosiphum padi* (L.)

**DOI:** 10.1101/2025.07.30.667415

**Authors:** Ilma A. Qonaah, Amma L. Simon, Duncan Warner, Toby J. A. Bruce, Rumiana V. Ray

## Abstract

**INTRODUCTION:** *Rhopalosiphum padi* (L.) is a vector of Barley Yellow Dwarf Virus (BYDV) infecting major cereals including wheat. Recently, a winter wheat variety (G1) was identified as exhibiting significant aphid resistance through antixenosis and antibiosis. This study compares resistance to viruliferous aphids, and BYDV transmission, in G1 with RGT Wolverine and RGT Illustrious, a BYDV resistant and susceptible wheat varieties, respectively. We aimed to define how aphid resistance affects BYDV transmission, infection and spread.

**RESULTS:** Seedling choice and olfactometer bioassays using wheat volatile organic compounds revealed that G1 emits an aphid repellent compound, identified as 2-tridecanone using GC-MS. Electrical penetration graph recordings showed restricted phloem access and salivation of viruliferous *R. padi* in G1, associated with lower BYDV transmission efficiency. Quantitative RT-PCR revealed a three-fold reduction in BYDV gene expression ratio on G1 transmission leaves compared to RGT Wolverine or RGT Illustrious. In contrast, reduced systemic infection in RGT Wolverine implied a BYDV resistance mechanism of limiting viral replication and/or movement. Rearing aphids on the aphid or BYDV resistant varieties modified their host selection behaviours suggesting vector conditioning with implications for viral transmission and spread on susceptible hosts.

**CONCLUSION:** This study demonstrates that the aphid resistance in G1 reduced BYDV transmission. Contrastingly, RGT Wolverine appeared to limit systemic viral infection despite high transmission efficiency. Combining these two distinct resistance mechanisms by breeding offers valuable strategy against both the aphid and the virus. To further define aphid and BYDV defence responses in G1, transcriptomic and metabolomic studies will be required.

## 1. Introduction

Barley Yellow Dwarf Virus (BYDV), in the Luteoviridae family, is a persistent virus causing disease in more than 150 plant species including economically important cereals such as wheat (*Triticum aestivum* L.)^1,2^. Alate *Rhopalosiphum padi* (L.) transmits BYDV-PAV in a circulative and non-propagative manner^3^, particularly in the autumn just after crop emergence, and in the spring during tillering^4,5^. BYDV infection is characterised by stunted growth and leaf chlorosis and is associated with up to 86% yield loss in wheat^6–8^.

The most sustainable strategy to control BYDV is by increasing host resistance to the vector and/or to the virus^9^. Recently, the wheat variety RGT Wolverine carrying *Bdv2* was commercially introduced in the United Kingdom^10^. *Bdv2* is a genetically dominant gene originating from the wild wheat relative *Thinopyron intermedium,* providing resistance against PAV, GAV, and GPV serotypes^11,12^. However, how *Bdv2* resistance in RGT Wolverine operates is unclear, with past literature speculating potential mechanisms including suppression of viral replication and movement or reduced transmission due to effects on aphid fitness^13,14^.

Aphid resistance in commercial wheat varieties has not yet been identified, but recently we found a variety, designated as G1, that exhibited antibiosis (feeding difficulties and reduced survival) and antixenosis (reduced alate attraction and settlement) with non-viruliferous *R. padi*^15,16^. Since BYDV infection is known to modify insect settlement and feeding behaviours with benefits for BYDV transmission and spread^17^, it is unclear if the identified resistance in G1 remains effective against viruliferous aphids. The objectives of this work were therefore to determine whether the previously identified aphid resistance in the G1 wheat genotype is active in the presence of BYDV in *R. padi* and to define how aphid resistance affects BYDV transmission, infection and spread. Furthermore, one volatile organic compound (VOC) from G1 headspace sample was identified through GC-MS, and its effects on aphid behaviour and fitness were evaluated to assess its contribution towards G1 resistance. We hypothesized that aphid resistance in G1 reduces BYDV transmission, characterised visually and by RT-PCR as viral gene expression. Aphid bioassays using the electrical penetration graph (EPG) technique were used to investigate aphid feeding behavioural traits relevant to viral transmission. Finally, the effects of BYDV and aphid resistance of the rearing wheat genotype on host selection by the aphid-vector were explored to determine if variation in host resistance impacted subsequent aphid behaviours and BYDV transmission.

### 2. Materials and methods

#### 2.1 Plants and insects

A panel of winter wheat (*Triticum aestivum* L.) genotypes previously screened for aphid resistance was provided by Syngenta Ltd. and assessed at The University of Nottingham, Sutton Bonington^16^. Due to commercially sensitive information, a unique code name consistent with previously published work on phenotyping these lines for aphid resistance was used for each variety, namely S3, S4, G1, and RGT Illustrious previously referred to as R1^16^. RGT Illustrious was included due to its susceptibility towards both *Sitobion avenae* (F.) and *Rhopalosiphum padi* (L.), characterised by high alate settlement and aphid attraction to its VOCs ^16^. S3 and S4 also presented antibiosis susceptibility, exhibited by long feeding duration and high aphid survival^16^. RGT Wolverine, with known BYDV resistance, was used as control. Wheat plants were sown using Tray Substrate Growcoon (Klassmann-Deilmann GmBH, Germany) in a 96 well potting tray (module size; 3.8 x 3.8 x 7.8 cm) in a glasshouse at 20 ± 2°C. Experiments were carried out using plants at 3-5 leaves seedling stage (GS15). For VOC collection at mid-flowering stage (GS65), plants were vernalised for 6 weeks at 4°C before being potted in John Innes No. 2 compost and grown under glasshouse conditions at 20 ± 2°C.

Barley Yellow Virus-PAV *Rhopalosiphum padi* (L.) were obtained from the National Institute of Agricultural Botany, Cambridge. Viruliferous aphids were fed on oat (cv. Gerald) rearing plants for 4 days, after which all aphids were removed. Then a subset of a non-viruliferous *R. padi* population, originally obtained from Rothamstead Research, Hertfordshire, also reared on oats (cv. Gerald), was introduced to create comparable (viruliferous and non-viruliferous) populations of the same *R. padi* clone. Non-viruliferous and viruliferous aphids were reared in separate Bugdorm-4 insect rearing cages (47.5 cm^3^, NHBS Ltd, UK) incubated under the same controlled environment conditions at 20°C day temperature, 18°C night temperature, 16:8 light:dark photoperiod.

#### 2.2 Aphid settlement, volatile collection and olfactometer assays

Eight replicates of G1, RGT Illustrious, S3, S4, and RGT Wolverine were randomly arranged in a Bugdorm-4 insect rearing cage (47.5 cm^3^) in potting trays (module size; 5 x 5 x 6 cm) for aphid settlement assays. Following one hour of starvation, 100 *R. padi* alates carrying BYDV were released into the cage from a falcon tube and the number of alates settled on each plant was recorded after four hours^16^. This experiment was carried out twice in a glasshouse maintained at 20 ± 2°C.

The VOC headspace samples used in this study were collected from uninfected mid-flowering (GS65) wheat plants using Pyetech volatile collection kit, following a previously described method^16,18^. The same VOC samples were also used in olfactometer behavioural assays in the previous publication^16^.

Behavioural responses of viruliferous alate *R. padi* towards wheat VOCs were assessed using the same method as described by Qonaah et al. (2024), with RGT Illustrious as control in a four-arm olfactometer and G1, S3, S4, or RGT Wolverine VOCs on the three arms^19,20^. The time alate spent in olfactometer zones were recorded for 16 minutes using olfactometeR package (https://rdrr.io/github/Dr-Joe-Roberts/olfactometeR/) on RStudio R 4.1.2 (RStudio, Massachusetts, USA), with 45° rotation every two minutes to avoid lighting bias and standardised room temperature at 25°C^21,22^. One experiment was carried out with ten replicates.

#### 2.3 Wheat volatile compound identification, quantification, and validation

The collected G1, RGT Illustrious, and RGT Wolverine volatiles were analysed using an Agilent 7820A GC with a 5977B single quad mass selective detector (Agilent Technologies, Cheadle, UK) fitted with a non-polar HP5-MS capillary column^23^. Automated injections of 1µl aliquots using a G4513A autosampler (Agilent Technologies) were made in splitless mode at 285°C. The oven program was 35°C for 5 minutes, followed by a gradual increase of 10°C min^-1^ to 285°C. Before and after plant VOC samples, a single run of *n-*alkanes and blank sample (conditioned Porapak Q eluted with 750µl DCM) was injected. Agilent MassHunter Quantitative Analysis with Quant-my-way interface was used to remove background noise on blank samples, retention time and peak area acquisition. Tentative compound identification was based on mass spectra, NIST library and *n*-alkanes linear retention times^24^. 2-Tridecanone was confirmed and quantified through co-chromatography with 100 ng μl^-1^ of authentic compound.

Behavioural responses of aphids to 2-tridecanone were tested using a four-arm olfactometer assay. 2-Tridecanone was diluted to 16 ng μl^-1^ as quantified during co-chromatography in diethyl ether^25^. The assay was carried out as described in 2.2, with three blank arms and one test compound arm, with one experiment made of ten replicates.

#### 2.4 Insecticidal activity assessment of wheat volatiles

2-Tridecanone, identified as the main significant compound (VIP > 1) present in the VOCs of G1 was validated and quantified via co-injection before being assessed for its effects on aphid life-history traits. The experiments were conducted in a glasshouse with temperature maintained at 20 ± 2°C. First, *R. padi* alates were exposed to 0.16 ng μl^-1^ 2-tridecanone using four-arm olfactometer for one hour, with the 10 μl of the compound placed in each of the four arms and replaced every 20 minutes. Three alates were placed on the main leaf of RGT Illustrious at GS15, secured with clip cage. A minimum of ten replicates for each viruliferous and non-viruliferous *R. padi* were included in the analysis, excluding a couple reps with dislodge clip cages. Alate survival and number of nymphs were recorded after seven days. The efficacy of 2-tridecanone as foliar treatment was also evaluated. To ensure the safety of user and environment, 10% ethanol was used as solvent to produce 0.16 ng μl^-1^ 2-tridecanone. In a Bugdorm-4 insect rearing cage (47.5 cm^3^, NHBS Ltd, UK), 12 biological replicates of GS15 stage G1 and RGT Illustrious seedlings were uniformly sprayed with 10 ml of 0.16 ng μl^-1^ 2-tridecanone or 10% ethanol as control. Three *R. padi* alates were placed on the main wheat leaf secured with a clip cage. Alate survival and number of nymphs produced were recorded after seven days.

#### 2.5 EPG feeding assay

To assess feeding behaviour post settlement, an electrical penetration graph (EPG) Giga 8-dd (EPG System, Wageningen, The Netherlands)^26^ was used and set up detailed in Qonaah et al. (2024). Wingless (apterous) viruliferous *R. padi* (5-7 days old) were used in this experiment. Aphids were lowered onto the main leaf of a wheat plant (GS15) once EPG reading was started. The experiment was conducted inside a Faraday cage with a standardised room temperature of 25°C. Data were collected using EPG Stylet+d software for eight hours and annotated using EPG Stylet+a software (EPG System, Wageningen, The Netherlands). Replicates with no feeding activity in the first hour or continuously for three hours at any point during the assay were discarded. At least 15 valid biological replicates were obtained for each variety.

#### 2.6 Antibiosis assay

Life history traits assessment of viruliferous *R. padi* was performed in a growth chamber with 20°C day temperature, 18°C night temperature, 16:8 light:dark photoperiod. In all three experiments conducted, *R. padi* (+BYDV) alates were placed on the main leaf of GS15 wheat, encased in clip cages as described by Qonaah et al. (2024). Alate aphid mortality and reproduction were recorded after seven days. Nymphs produced by the alates were placed back onto the leaf to record their mortality after seven days. For two experiments, one alate was placed on each plant at the start of the experiment, and one nymph placed on each plant on the subsequent week. During the third experiment three alates were placed on each replicate at the beginning, and three nymphs placed on each replicate on the following week. In total, there were three experiments with four replicates each.

#### 2.7 Viral transmission and systemic infection

To investigate the relationship between initial vector numbers, local and systemic BYDV infection, and to compare transmission in the different wheat varieties, the main leaf of RGT Illustrious, G1, and RGT Wolverine at GS15 was infested with 5, 10, or 25 viruliferous apterous *R. padi*. Aphids were allowed to feed for 24 hours to inoculate the plants with BYDV before being removed^27,28^. The inoculated main leaf was harvested immediately after aphid removal. To assess local viral transmission efficiency, one experiment was carried out with three replicates for each genotype and treatment (5, 10, or 25 viruliferous aphids). A second set of plants was used to assess systemic infection by harvesting distal, non-inoculated leaves, at 7 and 14 days post BYDV inoculation. One experiment with three replicates for each genotype and treatment (5, 10, or 25 viruliferous aphids) was caried out. The experiments were performed in a growth chamber with 20°C day temperature, 18°C night temperature, 16:8 light:dark photoperiod with harvested leaf material being used for RNA extraction for BYDV gene expression.

#### 2.8 BYDV assessment in different wheat varieties

BYDV symptom expression in different wheat varieties was evaluated following transmission of the virus by 10 apterous viruliferous *R. padi.* This number was shown to provide the most consistent local viral transmission across genotypes in the BYDV transmission experiments. Aphids were allowed to feed for 24 hours on a wheat leaf (GS15) before their removal^27^. Symptoms of BYDV were visually scored from 0 to 5 with score of 0 indicating no symptoms, 1 minimal chlorosis on a couple of leaves, 2 chlorosis on 20% of the plant, 3 for 40% chlorosis, 4 for 60% chlorosis and 5 more than 70% chlorosis of the plant^29^. Plants were assessed every 7 days for 21 day-period. At day 21, leaf material of three replicates was harvested for RNA extraction for BYDV gene expression using qPCR. The experiment was repeated twice with a minimum of ten replicates for each variety in a growth chamber with 20°C day temperature, 18°C night temperature, 16:8 light:dark photoperiod.

#### 2.9 BYDV gene expression

RNA extraction was carried out using Qiagen RNeasy plant mini kit, 500 ng of RNA was used for cDNA synthesis using iScript cDNA synthesis kit and subsequently diluted to 20 ng μl^-1^ concentration. Quantification of BYDV gene expression was performed using Bio-Rad CFX96 Touch Real-Time PCR System, with Bio-Rad SsoAdvanced Universal SYBR master mix, 5 μl cDNA in a total reaction volume of 13 μl, using BYDV-PAV primers listed by Balaji et al. (2003)^30^. Housekeeping genes used as controls were GADPH and TUBB^31^. Initial denaturation was set at 95°C for 10 minutes, with 40 cycles of 95°C denaturation for 15 seconds, 60°C annealing for one minute, 72°C extension for 10 seconds, and gradual increase to 95°C at 0.5°C min^-1^ on the final cycle. Gene expression of BYDV was measured as comparative ratio 2^-ΔCq^ [ΔCq = Cq_BYDV_ – Cq_HKG_] against housekeeping genes (HKG), regularly applied to quantify viral expression^32–35^. The average Cq of GADPH and TUBB was used as HKG.

#### 2.10 Effect of rearing wheat genotype on aphid behaviour and viral transmission

To determine whether BYDV and aphid resistance influence on vector behaviour, *R. padi* settlement assays were carried out after the aphids were reared on G1, RGT Illustrious, and RGT Wolverine. In a growth chamber, with 20°C day temperature, 18°C night temperature, 16:8 light:dark photoperiod, the wheat varieties G1, RGT Illustrious, and RGT Wolverine, at GS15 stage, were covered in individual nets and 20 alate *R. padi* were placed on the plant. After 2 weeks, the aphid infested plant was placed in the centre of BugDorm-4 insect rearing cage (47.5 cm^3^) and clean G1, RGT Illustrious, and RGT Wolverine (GS15) were introduced into the cage being placed equidistant (20 cm) from the infested host plants. The number of alates and apterous aphids settling on introduced plants was recorded after 2, 4, 6, 24, and 48 hours. The total aphid weight on the rearing plant was recorded after the experiment. Two experiments with four replicates for viruliferous or non-viruliferous aphids each were conducted. BYDV symptoms were recorded and in the second experiment plant material from the rearing and introduced plant with viruliferous aphids were collected for BYDV gene quantification.

#### 2.11 Statistical analysis

Statistical analysis was carried out using Genstat 23^rd^ (VSN International, Hemel Hempstead, UK). Generalised linear model (GLM) with binomial distribution and link function logit was used to analyse proportional data including alate settlement, survival, and BYDV visual symptom score. Analysis of continuous data including duration of probing activities during EPG, and aphid weight were carried out using GLM with link function square root or log. Comparative ratio Cq values were analysed using GLM with poisson distribution and link function log, like other count data including probes during EPG. Within the GLM, experiment and replicate factors are included, if significant, to the model. Interaction with time was included to analyse stylet derailment period during EPG and BYDV gene expression in distal leaf. Pairwise comparison was done with Fisher’s LSD. Analysis of number of entries into and time aphid spent in an olfactometer zone were completed using T-test or Mann-Whitney test for parametric and non-parametric data, respectively. EPG data was analysed using RStudio 4.2.3 (Rstudio, Massachuchetts, USA). Non-linear data obtained during EPG, namely number of salivation events and feeding and salivation time, were assessed using asymptotic nonlinear regression from aomisc package on RStudio^36^.

Simca P+13 software (Umetrics, Umeå, Sweden) was utilised for volatile profiling analysis and visualization. Orthogonal partial least squares-discrimination analysis (OPLS-DA) was performed using the peak areas of detected volatiles as dependent variables. The robustness of the OPLS-DA was determined through the model fitness (R^2^) and predictive ability (Q2) values, as detailed in Table S1. Compounds with a variable importance in the projection (VIP) score of >1 were considered significantly different.

## 3. Results

### 3.1 Pre alighting cues: plant settlement and olfactometer bioassay

Behaviour of viruliferous alate *R. padi* was observed separately in plant settlement bioassays on whole wheat seedlings or in an olfactometer in the presence of collected VOCs (Fig. 1). Plant settlement bioassays revealed clear aphid preference for RGT Illustrious because viruliferous alate *R. padi* settled four times more on RGT Illustrious than on G1, RGT Wolverine, and S3 (Fig. 1a).

**Figure 1.**
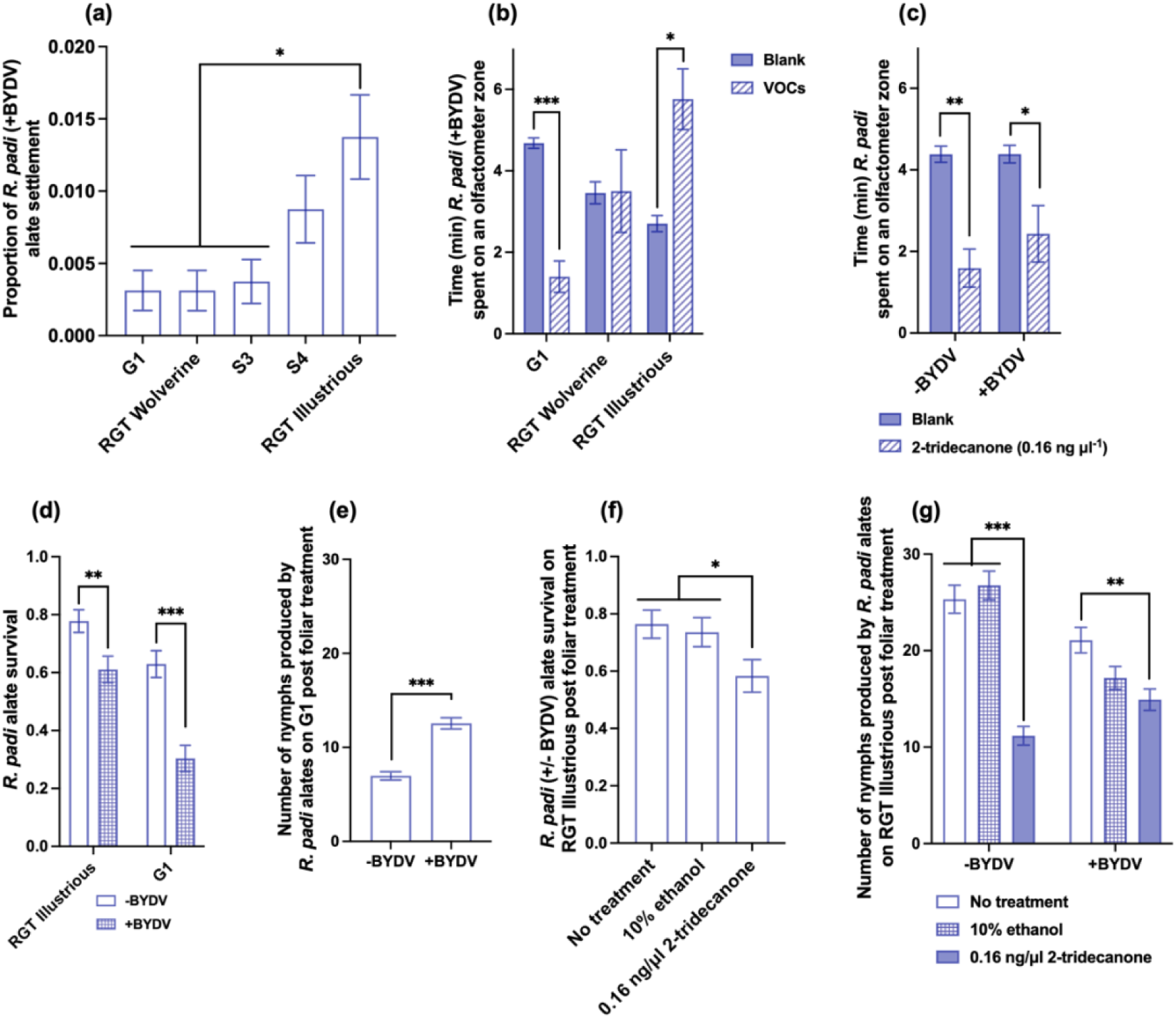
Pre-alighting behaviour of BYDV-PAV alate *R. padi* and effects of 2-tridecanone volatile organic compound (VOC) on aphid behaviour and as foliar treatment. (a) Alate settlement on whole wheat seedlings analysed using generalised linear model (GLM) with binomial distribution (*P* < 0.001, *n* = 16). Time alate spent on (b) wheat headspace VOCs against blank and (c) on 2-tridecanone against blanks, analysed using T-test and Mann-Whitney test (*n* = 10). Alate survival and number of nymphs were analysed using GLM with binomial and poisson distribution, respectively. (d) Alates with BYDV exhibited lower survival than nonviruliferous alates both on RGT Illustrious (*P* = 0.007, *n* = 36) and G1 (*P* < 0.001, *n* = 36), regardless of foliar treatments. (e) Viruliferous alates on G1 produced higher (*P* < 0.001) number of nymphs than nonviruliferous alates, regardless of foliar treatments (*n* = 36). (f) Alate survival of both viruliferous and non-viruliferous *R. padi* on RGT Illustrious exhibited significantly (*P* = 0.043) reduced survival on 2-tridecanone treated plants (*n* = 24). (g) Foliar treatment of 2-tridecanone on RGT Illustrious significantly reduced number of nymphs for nonviruliferous *R. padi* (*P* < 0.001, *n* = 12). * *P* ≤ 0.05; ** *P* ≤ 0.01; *** *P* ≤ 0.001

Olfactometer bioassays showed aphids spent significantly more time (approximately twice as much time) in the presence of RGT Illustrious VOCs compared to RGT Wolverine, G1, and S4 VOCs (Fig. S1a). To confirm behavioural responses, the VOCs of RGT Wolverine, G1 and RGT Illustrious were further tested on olfactometer against blanks. These varieties were chosen because they had the most significant differences in the previous experiment. Viruliferous *R. padi* spent four times longer in zones with the blank than with G1 VOCs, exhibiting repulsion (Fig. 1b). In contrast, the aphids displayed clear attraction towards RGT Illustrious, spending 100% more time on RGT Illustrious VOCs compared to blanks, while RGT Wolverine VOCs did not elicit any significant response (Fig. 1b).

### 3.2 Wheat VOCs compound identification, quantification, and validation

Putative bioactive compounds of VOCs collected at flowering of G1, RGT Illustrious, and RGT Wolverine were identified following GC-MS analysis of headspace samples. Out of 102 peaks obtained, OPLS-DA analysis showed 37 statistically significant (VIP > 1) peaks between the samples, with 36 unknown peaks. One compound, 2-tridecanone, was identified and quantified at 0.16 ng μl^-1^ only from G1 VOCs (Table S1-S2, Fig. S1b). This compound was specific for G1 as was not found in VOCs of RGT Illustrious or RGT Wolverine. When tested in olfactometer bioassays, both non-viruliferous and viruliferous *R. padi* spent significantly less time in the 2-tridecanone arm compared to the blank (Fig. 1c).

### 3.3 Effects of 2-tridecanone on *R. padi* alate survival and reproduction

To determine if 2-tridecanone impacted aphid life history traits, 2-tridecanone was applied as priming agent on *R. padi* alates and as foliar treatment on resistant G1 and susceptible RGT Illustrious plants. Chemically priming aphids led to reduced alate survival and number of nymphs produced, regardless of BYDV presence (Fig. S1c-d).

Foliar treatment of resistant G1 and susceptible RGT Illustrious leaves with 2-tridecanone was performed to evaluate the potential effects of the compound against aphids on their plant hosts (Fig. 1d-g). Overall, *R. padi* alates carrying BYDV displayed lower alate survival on RGT Illustrious and G1 (Fig. 1d), though alates with BYDV produced 100% more nymphs than nonviruliferous alates on G1 (Fig 1e). Regardless of BYDV presence, foliar application of 0.16 ng μl^-1^ of 2-tridecanone on the susceptible RGT Illustrious led to a 22% reduction in survival of *R. padi* alates (Fig. 1f) also decreasing nymph production. 2-tridecanone treatment of the aphid susceptible RGT Illustrious resulted in 100% reduction in non-viruliferous *R. padi* nymph production and fewer nymphs of viruliferous *R. padi* (Fig. 1g). However, 2-tridecanone failed to affect alate survival and reproduction on G1 (Fig. S1e). There were no differences in BYDV symptoms following foliar treatment of plants with 2-tridecanone (Fig S1f).

### 3.4 Post-alighting cues: feeding behaviour and life-history traits

EPG assay was used to observe differences in feeding activity of viruliferous *R. padi* on susceptible, BYDV resistant, and aphid resistant plants. Upon settling, viruliferous *R. padi* exhibited the shortest first probing period on RGT Wolverine at 35.32 minutes, followed by S4 and RGT Illustrious (Fig. 2a). As the aphid’s stylet enters the mesophyll, it may derail, and this was found to occur most frequently on G1 (Fig. 2b). There was a significantly longer derailment time on G1 than the other varieties five hours into probing, rising steeply to 172.77 minutes after eight hours, more than double the derailment time observed on other varieties (Fig. 2c).

**Figure 2.**
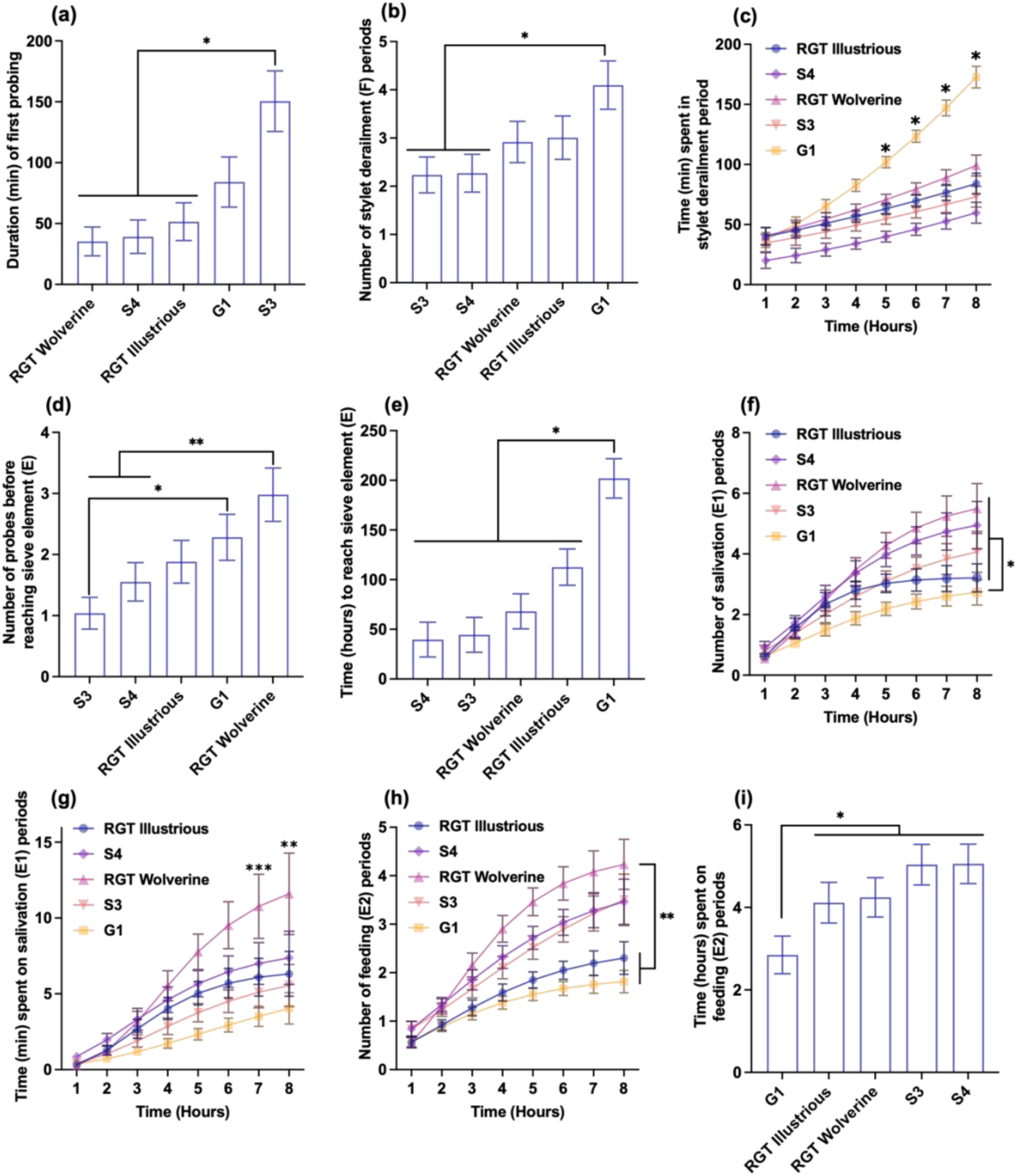
Electrical penetration graph (EPG) reading of *R. padi* with BYDV feeding behaviour (*n* = 15-17). (a) Duration of first probing analysed using generalised linear model (GLM) with normal distribution and link function log (*P* < 0.001). (b) Number of stylet derailment analysed using GLM with poisson distribution and link function log (*P* = 0.003). (c) Time spent in stylet derailment analysed using GLM with normal distribution and link function square root (*P* < 0.001). (d) number of probes before sieve element analysed using GLM with poisson distribution and link function log (*P* < 0.001). (e) Time aphid took to reach sieve element analysed using GLM with normal distribution and link function log (*P* < 0.001). (f) Number and (g) time of salivation periods analysed using asymptotic nonlinear regression (*P* < 0.001). (h) Number of feeding periods analysed using asymptotic nonlinear regression (*P* < 0.001). (i) Time of feeding period analysed using GLM with normal distribution and link function log (*P* = 0.009). * *P* ≤ 0.05; ** *P* ≤ 0.01; *** *P* ≤ 0.001

Number of probes before the stylet reached sieve element was also high on G1, second to RGT Wolverine (Fig. 2d), with the aphid taking the longest time to reach sieve element of G1 at 201.95 minutes, five times longer than on S4 (Fig. 2e). Once reaching the sieve element, aphids displayed the highest salivation frequency on RGT Wolverine whilst frequency on G1 was the lowest of all varieties (Fig. 2f). After seven hours, the time aphids spent salivating was highest on RGT Wolverine at 10.77 minutes and lowest on G1 at 3.52 minutes (Fig. 2g). Feeding frequency followed a similar trend and was significantly more frequent on RGT Wolverine than on G1 or RGT Illustrious (Fig. 2h). Overall, viruliferous *R. padi* spent less time feeding on G1 at 2.8 hours, 33% less than on RGT Illustrious and RGT Wolverine, and 45% less than on S3 and S4 (Fig. 2i).

To investigate the fitness of viruliferous *R. padi* alates and their progeny in the event of alate settlement, clip cages were used to confine aphids on a plant leaves. Alate survival, number of nymphs produced, and nymph survival were highest on RGT Illustrious, significantly different to G1 in all parameters (Fig. S2a-c).

### 3.5 Viral transmission and systemic infection

BYDV transmission and systemic infection were investigated on RGT Illustrious (aphid susceptible variety) and G1 (aphid resistant variety) alongside RGT Wolverine (known BYDV resistant variety) as a control. Viral gene expression ratio increased proportionally to the number of aphids used for inoculation: 25 aphids for inoculation transmitted three times more virus than 10 aphids, and eight times more than 5 aphids (Fig. 3a). Irrespective of initial vector numbers, BYDV expression ratio was significantly lower on G1 compared to RGT Illustrious and RGT Wolverine, the latter exhibiting three-fold higher expression ratio relative to G1 (Fig. 3b). In contrast, systemic infection measured here by viral expression on the distal leaves showed the lowest BYDV expression ratio on RGT Wolverine at 7- and 14-days post inoculation (Fig. 3c-e). In G1, systemic infection was the highest out of all three varieties tested at 7 dpi but was reduced at 14 dpi. At 14 dpi G1 and RGT Illustrious had similar systemic infection levels but RGT Wolverine was still much lower (Fig. 3c-e).

**Figure 3.**
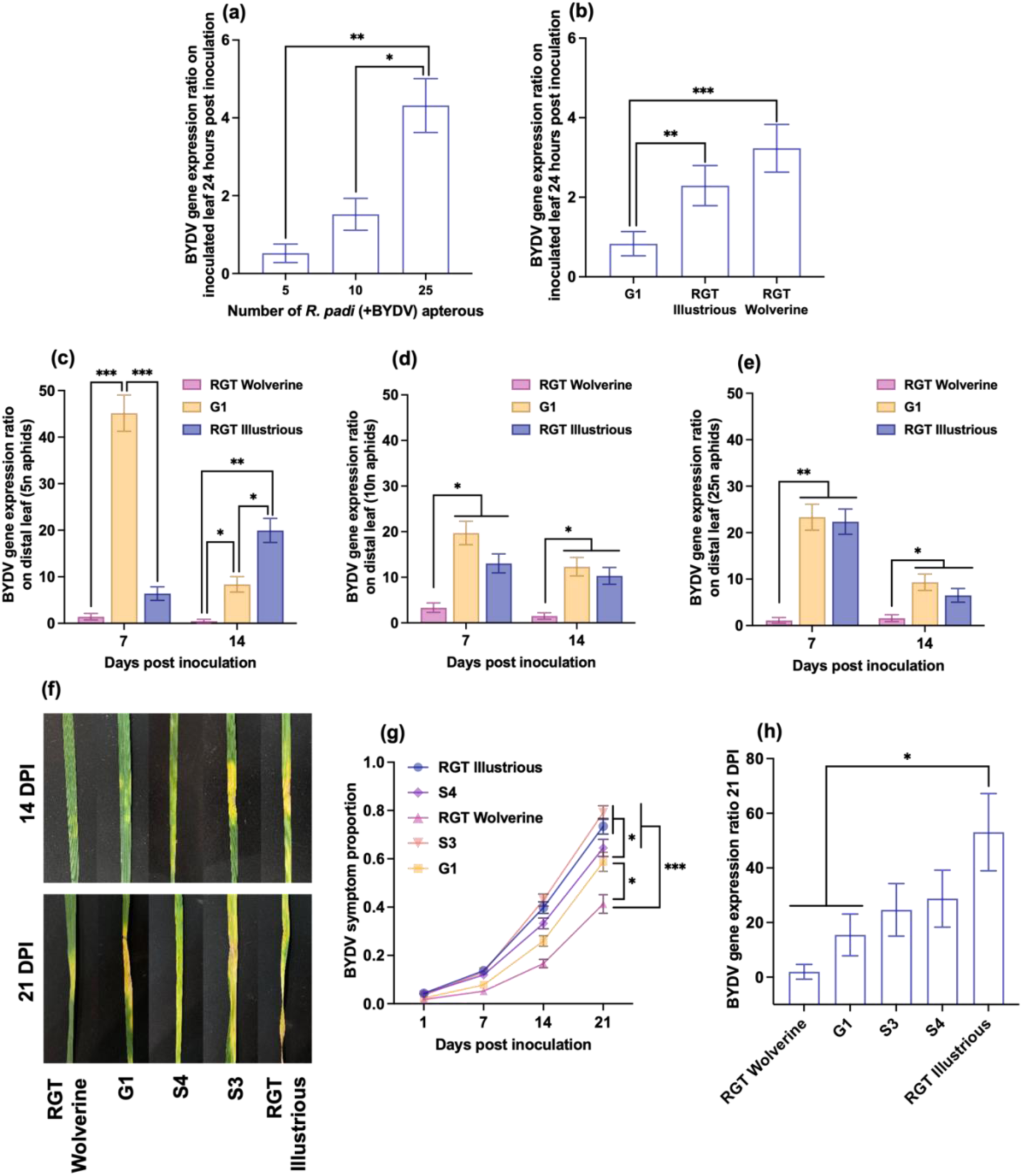
BYDV symptoms and gene expression. BYDV gene expression ratio analysed using generalised linear model (GLM) with poisson distribution and link function log on (a-b) inoculated leaf 24 hours post inoculation (*P* < 0.001, *n* = 9), and distal leaf of plants inoculated with (c) 5, (d) 10, and (e) 25 *R. padi* apterous 7 and 14 dpi (*P* < 0.001, *n* = 3). (f-g) BYDV symptoms post inoculation with 10 apterous *R. padi* for 24 hours analysed using GLM with binomial distribution and link function logit (*P* < 0.001, *n* = 24). (h) BYDV gene expression ratio whole plant 21 days post inoculation (*P* < 0.001, *n* = 6) analysed using GLM with poisson distribution and link function log * *P* ≤ 0.05; ** *P* ≤ 0.01; *** *P* ≤ 0.001

Marked reduction in BYDV gene expression ratio of 80% was observed 14 dpi compared to 7 dpi on plants inoculated with 5 aphids (Fig. 3c). A similar pattern was observed following inoculation with 10 or 25 aphids (Fig. 3d-e). In contrast, BYDV expression ratio on distal RGT Illustrious leaves was 68% higher at 14 dpi compared to 7 dpi after inoculation with 5 aphids (Fig. 3c), with no change observed with 10 aphids (Fig. 3d), and a decrease with 25 aphids (Fig. 3e).

After 21 days, BYDV symptoms were significantly less severe on RGT Wolverine, than other varieties, followed by G1 with 20% less severe symptoms than S3 or RGT Illustrious (Fig. 3f-g). BYDV gene expression was lowest on RGT Wolverine, in contrast to RGT Illustrious showing significantly higher viral expression than both G1 and RGT Wolverine (Fig. 3h).

### 3.6 Effect of aphid and BYDV resistance on aphid behaviour and viral transmission

Alate and apterous aphid behaviour and BYDV expression were assessed after the alates were reared on G1, RGT Illustrious, and RGT Wolverine to investigate if variation in aphid and viral resistance of the primary host on which aphids are reared on affects aphid host preference and BYDV transmission.

Non-viruliferous alate *R. padi* reared on the BYDV susceptible RGT Illustrious or G1 displayed settlement preference for RGT Illustrious, from 4 hours post introduction (Fig. 4a-b). Non-viruliferous alates reared on RGT Wolverine displayed preference to RGT Illustrious and RGT Wolverine (Fig. 4c). The same effect was more noticeable for aphids carrying BYDV, with preferences for RGT Illustrious displayed by aphids reared on RGT Illustrious and G1 as early as 2 hours post introduction (Fig 4d-e). Viruliferous alates reared on RGT Wolverine preferred the introduced G1 or RGT Illustrious but avoided the RGT Wolverine (Fig. 4f). Overall, aphid population growth was the lowest when reared on G1, with 40% less total aphid weight compared to RGT Wolverine and RGT Illustrious, regardless of BYDV presence (Fig. 4g). RGT Wolverine as rearing host expressed the lowest BYDV symptom severity and gene expression ratio whilst RGT Illustrious was the highest, expressing 5- and 48-fold higher BYDV expression ratio relative to G1 and RGT Wolverine, respectively (Fig. 4h-i).

**Figure 4.**
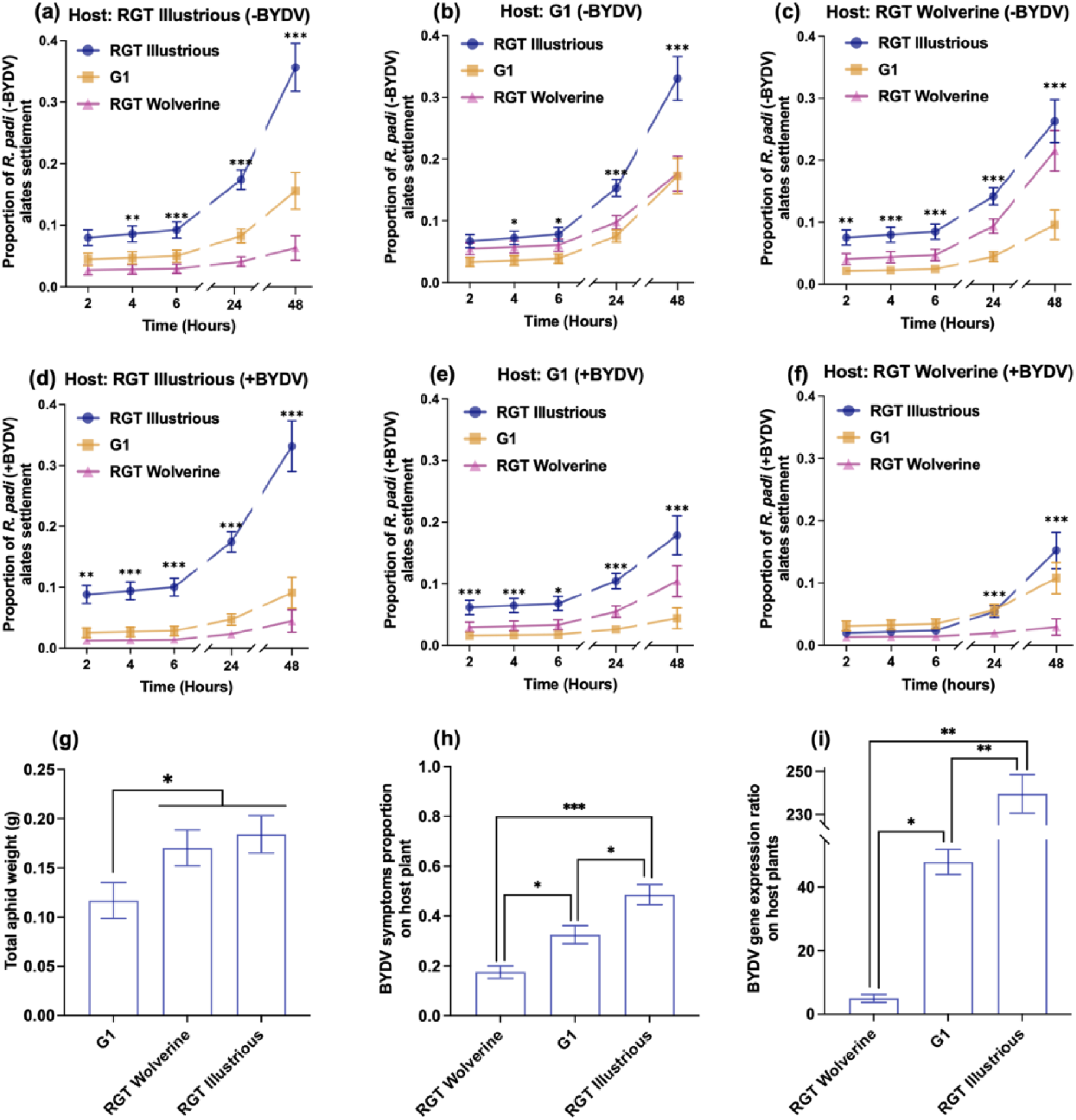
Effects of rearing genotype on alate *R. padi* host choice and initial BYDV infection, analysed using generalised linear model (GLM). Proportion of alate settlement of (a-c) nonviruliferous *R. padi* and (d-f) *R. padi* with BYDV for aphids reared on (a,d) G1, (b,e) RGT Illustrious, and (c,f) RGT Wolverine analysed with binomial distribution logit (*P* < 0.001, *n* = 8). (g) Total weight of *R. padi* reared on G1, RGT Illustrious, and RGT Wolverine for 14 days analysed with normal distribution (*P* < 0.001, *n* = 16). (h) BYDV symptoms on host plants analysed using binomial regression and link function logit (*P* = 0.015, *n* = 8). (i) BYDV gene expression on host plant analysed using poisson distribution and link function log (*P* < 0.001, *n* = 3). * *P* ≤ 0.05; ** *P* ≤ 0.01; *** *P* ≤ 0.001

Settlement of nonviruliferous *R. padi* apterae was uniform regardless of the rearing genotype, with 50-100% more aphids on RGT Illustrious than G1 and RGT Wolverine after 48 hours (Fig. 5a-c). There was a higher number of viruliferous *R. padi* apterous on the introduced RGT Illustrious, regardless of host genotype (Fig. 5d-f). Aphids reared on RGT Illustrious and RGT Wolverine selected RGT Wolverine least (Fig. 5d, f), while aphids reared on G1 were indifferent to the introduced G1 and RGT Wolverine (Fig. 5e).

**Figure 5.**
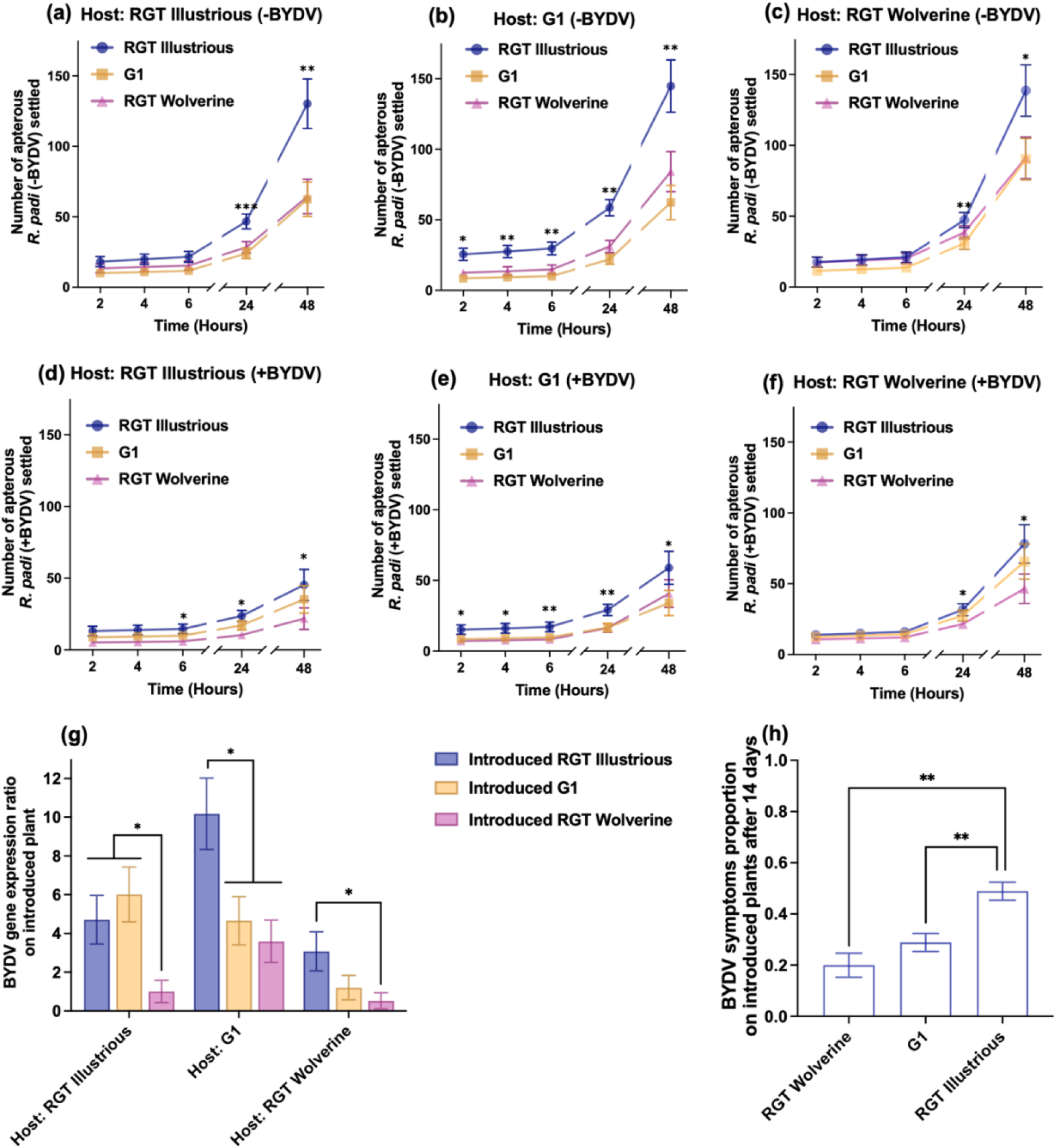
Effects of rearing genotype on non-viruliferous and BYDV infected apterous *R. padi* host choice and BYDV spread and transmission, analysed using generalised linear model (GLM). Number of (a-c) nonviruliferous *R. padi* and settled on introduced plants for aphids reared on (a) RGT Illustrious, (b) G1, and (c) RGT Wolverine analysed using poisson distribution and link punction log (*P* < 0.001). Number of viruliferous *R. padi* (d-f) settled on introduced plants for aphids reared on (d) RGT Illustrious, (e) G1, and (f) RGT Wolverine, analysed using poisson distribution and link function log (*P* < 0.001, *n* = 8). (g) BYDV gene expression ratio on introduced plants, analysed using poisson distribution and link function log (*P* < 0.001, *n* = 3). (h) BYDV symptoms on introduced plants two weeks after inoculation, analysed using binomial distribution and link function logit (*P* = 0.021, *n* = 9). * *P* ≤ 0.05; ** *P* ≤ 0.01; *** *P* ≤ 0.001

Viruliferous aphids reared on RGT Illustrious had similar BYDV expression ratio on introduced RGT Illustrious and G1 with lowest BYDV expression in introduced RGT Wolverine (Fig. 5g). Aphids reared on G1 caused the highest BYDV expression ratio in introduced RGT Illustrious, whilst introduced G1 and RGT Wolverine exhibited significantly lower BYDV expression (Fig. 5g). Aphids reared on RGT Wolverine transmitted lower BYDV overall, although highest expression ratio was still observed on the BYDV susceptible RGT Illustrious (Fig. 5g). Two weeks post inoculation, BYDV symptoms on introduced RGT Wolverine and RGT Illustrious were significantly less severe than symptoms on RGT Illustrious, regardless of rearing host (Fig. 5h).

## 4. Discussion

This study addresses the knowledge gap in the potential use of aphid-mediated resistance for mitigating BYDV transmission and spread. Aphid resistance in G1, expressed as antixenosis via repellent volatiles and antibiosis shown as suppressed sieve element access, resulted in reduced aphid survival, fecundity and BYDV transmission. This resistance is effective against initial BYDV infection by alate aphids, though it may amplify short-range BYDV transmission through increased movement of apterous aphids, particularly towards susceptible genotypes like RGT Illustrious^37^.These findings suggest that BYDV management in terms of initial infection by alates could be improved by using wheat varieties such as G1 that are resistant to the aphid vector.

Host selection by alate aphids play a pivotal role in BYDV transmission and spread as primary infection occurs during flight season in autumn when alates carry BYDV from previous crops and perennial grasses as they migrate into seedling and tillering wheat^6,8,38^. Here we observed alate *R. padi* carrying BYDV-PAV to demonstrate low settlement on seedlings of G1, RGT Wolverine, and S3, with significantly more aphids settling on the susceptible variety RGT Illustrious. This behaviour was identical to preference choices of nonviruliferous aphids reported in our prior study^16^. Further assessment using wheat VOCs suggested alate attraction towards RGT Illustrious and repulsion to G1. One repellent compound, 2-tridecanone, was identified from the G1 VOCs using GC-MS and validated in behavioural bioassays. Previously reported in the headspace sample of healthy flowering wheat^39,40^, aphid-infested wheat seedlings^41^, and wild tomato^42^, 2-tridecanone has been shown to deter *R. padi*^41,43^, corroborating our finding.

We found that pre-exposure of aphids to 2-tridecanone or treatment of plants with this volatile applied to the foliage, significantly reduced *R. padi* survival and nymph production in the susceptible RGT Illustrious. These effects were not large, suggesting that perhaps a higher concentration might be more effective. The effects of low concentration used here imply a plant defence inducing signalling effect rather than a direct toxic effect on aphids. This is in agreement with results on G1 plants, where 2-tridecanone was initially identified, where no additional reduction in aphid performance was observed. Furthermore, prior studies have reported 2-tridecanone inducing plant defence against pests such as orange wheat blossom midge ^39^ and *Aphis gossypii*^42,44^, reinforcing its potential for pest management. Whilst the mechanism remains unclear, it may relate to the regulation of cytochrome P450 genes, as observed on cotton bollworm^45,46^. Supplementing wheat at early emergence with 2-tridecanone could potentially enhance aphid and BYDV control within integrated pest management (IPM) strategies, with further transcriptomic studies needed on *R. padi* to better understand its effects.

In addition to antixenosis by repellent VOCs, aphid resistance in G1 is also expressed through restricted access to sieve elements, characterised by extensive stylet derailment, short salivation and phloem feeding by nonviruliferous aphids^16^. This trait is of interest because transmission of BYDV transpires most efficiently during aphid sustained salivation into the phloem^47^. Feeding behaviour of BYDV-infected *R. padi* on G1 was observed to align with patterns observed on nonviruliferous aphids. Extensive stylet derailment and time spent to reach sieve element indicated difficulties by aphids reaching the sieve element in G1 in addition to less salivation and short feeding periods. This contrasted with studies showing viral modifications of insect feeding behaviours to amplify transmission^48–50^ suggesting that the trait in G1 is constitutively expressed. This is consistent with observations that aphid resistance in G1, in the form of suppressed sieve element access, was not negated by BYDV presence suggesting that it in fact plays a role in BYDV resistance in the form of reduced virus transmission because it acts on the vector. In contrast, high salivation and phloem feeding period by viruliferous *R. padi* was observed on the BYDV resistant RGT Wolverine, indicating BYDV resistance in this genotype does not include inhibition of virus transmission by aphids.

Reduced BYDV transmission in G1, compared to other varieties, was indicated as its inoculated leaves expressed the lowest BYDV gene expression ratio in comparison to RGT Illustrious and RGT Wolverine. High BYDV gene expression ratio on RGT Wolverine was comparable to that of susceptible variety RGT Illustrious and was consistent with prolonged *R. padi* salivation time on the variety, confirming that transmission inhibition was not part of the viral resistance mechanism. BYDV expression on distal leaves of RGT Wolverine was the lowest, indicating suppression of viral replication and/or movement as the putative resistance mechanism as previously described^13^. A similar outcome is expected on adult stage wheat plants, as wheat carrying *Bdv2* gene has been reported to suppress BYDV replication and movement in both seedling and adult stage^14^.

Despite having a small brain, insects have been documented to have memory and learning ability for the sake of foraging, mating, and overall survival^51,52^. For instance, the model insect *Drosophila melanogaster* was able to display aversion towards an olfactory signal associated with electric shock^53^. Here associative learning in *R. padi* was demonstrated, evidenced by how the rearing host affected host selection. Both viruliferous and nonviruliferous *R. padi* reared on G1 exhibited preference towards other than G1 genotypes, possibly associating the pre-alighting cues of G1 with food scarcity, demonstrating aversive memory^53^. Meanwhile, *R. padi* reared on RGT Illustrious associated the pre-alighting cues of this genotype with food availability, amplifying RGT Illustrious preference and demonstrating appetitive memory^53^. Aphids reared on RGT Wolverine displayed more complex decision making. The nonviruliferous *R. padi* did not exhibit definitive aversion or attraction towards introduced RGT Wolverine. However, *R. padi* carrying BYDV avoided introduced RGT Wolverine, leading to comparable settlement on G1 and RGT Illustrious which was not observed in other treatments, likely a result of vector manipulation of aphid memory cues to ensure viral spread^54^.

The complexity of plant-aphid-virus interactions remains the main challenge in predicting disease outbreaks, particularly when incorporating emerging resistance traits. As a phloem-limited, persistently transmitted virus, BYDV relies on prolonged salivation and feeding for successful viral inoculation and acquisition^27,55^, which could lead to selection pressure for viral manipulation of the aphid host for increased reproduction. Interestingly, we observed a reduction in fecundity of viruliferous *R. padi* compared to non-viruliferous, contradicting this hypothesis but following previous findings^56–58^, suggesting BYDV infection to have an adverse effect on *R. padi* fitness.

High aphid settling bias and reproduction are expected to corelate with high BYDV disease incidence^59^. Aphid growth and BYDV disease on RGT Illustrious and G1 as the rearing host followed this model while RGT Wolverine did not. Despite high salivation and BYDV transmission on RGT Wolverine, suppressed systemic infection led to reduced viral inoculum available for acquisition by aphid progeny. It is likely that viruliferous aphids were able to detect this resistance mechanism during brief probing, leading to avoidance of RGT Wolverine observed during initial seedling choice assay, which was not caused by its VOCs.

On the aphid resistant G1, antibiosis suppressed aphid population growth and reduced BYDV transmission compared to the susceptible RGT Illustrious. This confirms that G1 antixenosis and antibiosis to the aphid vector can limit initial BYDV infection. However, these traits may inadvertently promote movement of virus-carrying aphids spreading BYDV to nearby plants^38^. The highest BYDV expression were observed in plants exposed to aphids from G1, suggesting that G1 does not prevent viral acquisition and infection by apterous individuals. This suggests that apterous aphids can potentially contribute to secondary spread and infection of neighbouring plants. Two weeks post-exposure, BYDV symptom severity on introduced plants followed the same pattern as on the original host, with most severity on RGT Illustrious, followed by G1, and least severe symptoms on RGT Wolverine, indicating that resistance traits of the rearing host do not alter long-term BYDV severity once the virus is established.

Data shown in this paper suggest that there are two aphid resistance mechanisms in G1: 1. emission of aphid repellent VOCs (during pre-alighting stage) and 2. suppression of sieve element access (during post-alighting stage). The latter is likely to result in aphids re-taking flight until they find more suitable hosts. Hence, both mechanisms resulted in low *R. padi* settlement on G1, regardless of BYDV presence. Most importantly, limited sieve element access in G1 reduced BYDV transmission, demonstrating a different BYDV resistance mechanism to *Bdv2* in RGT Wolverine. Aphid resistance in G1 is effective in limiting initial infection by BYDV but the lack of strong resistance against systemic infection is likely to result in field grown plants of this variety being sources for BYDV with G1-antixenosis favouring increased transmission to aphid-attractive but susceptible genotypes such as RGT Illustrious. Hence, the resistance is likely most beneficial in monoculture of the variety, where possibilities of virus transmission to other varieties are minimised or in mixtures where neighbouring plants have resistance to BYDV, such as RGT Wolverine. In contrast, RGT Wolverine was able to maintain low viral expression in distal to infection leaves indicating that replication and/or movement of BYDV within the host were impaired. This also meant that less inoculum was available to vectors to spread to new hosts. Combining the two distinct resistances in G1 and RGT Wolverine can lead to a potent new variety with capacity to reduce both transmission and spread of BYDV.

Further research, including transcriptomic and metabolomic analysis, is necessary to better understand aphid resistance in G1 and its implications for BYDV. Investigation of changes in wheat VOCs profile when infected with aphid and BYDV will also provide better understanding of the defence in each genotype. Furthermore, crossing populations of G1 and RGT Illustrious would be required to identify quantitative trait locus (QTLs) linked to the traits. Existence of an alternative BYDV resistance mechanism, associated with disruption of the aphid vector, could help as part of a resistance management strategy to reduce selection pressure for counter-resistance in BYDV to RGT Wolverine and other wheat varieties with the same resistance mechanism.

## Statements and declarations

### Data availability

The datasets generated during and/or analysed during the current study are available from the corresponding author on reasonable request.

## Acknowledgement

We would like to thank Samuel Asamoah for his assistance in wheat VOCs collection.

## Author contribution

IAQ designed and performed the experiments, analysed the data, and wrote the manuscript. ALS assisted experiment design and data analysis for olfactometer assay and EPG feeding assay. DW and TJAB reviewed and commented on the manuscript. RVR acted as the main advisor for the work and assisted in writing the manuscript. All authors contributed, read, and approved the manuscript.

