## Supplementary files for "An aphid resistant wheat variety reduces the transmission of Barley Yellow Dwarf Virus (BYDV) by *Rhopalosiphum padi* (L.)"

**Table S1**. Model fitness (R^2^) and predictive ability (Q2) values of Partial least squares-discrimination analysis (PLS-DA) for comparisons of G1, RGT Illustrious, and RGT Wolverine headspace volatile profiles.

| **Variety comparison** | **R^2^X(cum)** | **R^2^Y(cum)** | **Q2(cum)** |
| --- | --- | --- | --- |
| G1 vs RGT Illustrious | 0.82 | 1 | 0.999 |
| G1 vs RGT Wolverine | 1 | 1 | 1 |
| RGT Illustrious vs RG Wolverine | 0.83 | 1 | 0.998 |

**Table S2**. Putative compound identified from VOCs of G1 using GC-MS, validated and quantified through co-injection with synthetic compound obtained from ThermoFisher.

| **Putative**  **compound** | **Genotype** | **Presence** | **Abundance (ng μl^-1^)** |
| --- | --- | --- | --- |
| 2-Tridecanone | G1 | ✔︎ | 0.16 |
|  | RGT Illustrious | ✖︎ | - |

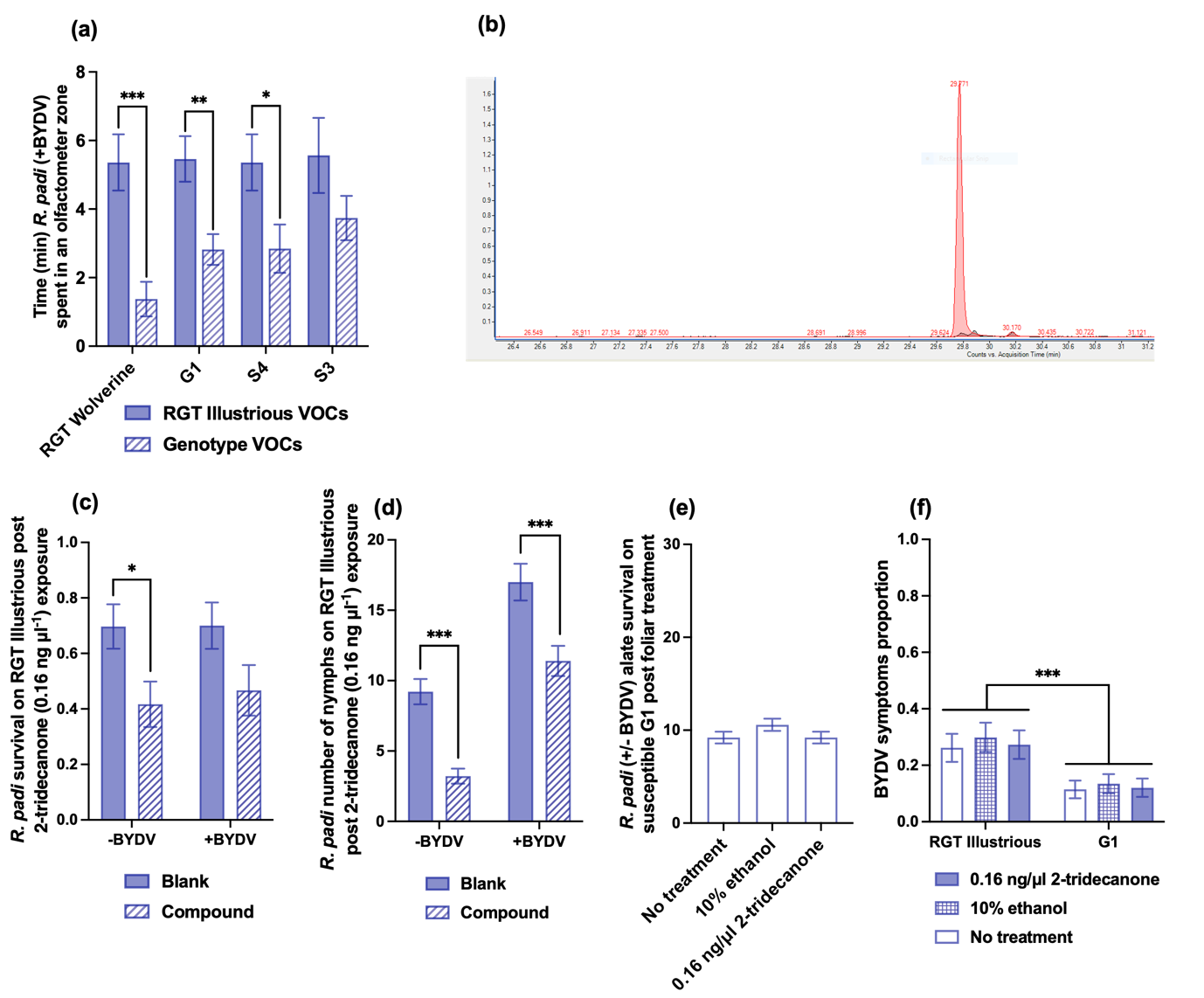

**Figure S1**. Viruliferous *R. padi* behaviour, volatile organic (VOC) compound detection, and assessment of 2-tridecanone volatile compound effects. (a) Time alate spent in an olfactometer zones with RGT Illustrious VOCs as control, analysed using T-test and Mann-Whitney test (*n* = 10). (b) 2-Tridecanone peak confirmation using GC-MS co-injection. (c) *R. padi* alates survival on RGT Illustrious following 0.16 ng μl^-1^ 2-tridecanone exposure, analysed using generalised linear model (GLM) with binomial regression and link function logit (*n* = 10-12, *P* = 0.018). (d) Number of nymphs produced by *R. padi* alates on RGT Illustrious following ng μl^-1^ 2-tridecanone exposure, analysed using GLM with poisson distribution and link function log (*n* = 10-12, *P* < 0.001). (e) Number of nymphs produced by R. padi alates on G1 following foliar treatments, analysed using GLM with poisson distribution and link function log (*n* = 12, *P* = 0.17). (f) Proportion of BYDV symptoms 14 days after foliar treatment and placement of *R. padi* carrying BYDV-PAV, analysed using generalised linear model (GLM) with binomial regression and link function logit (*n* = 12) * *P* ≤ 0.05; ** *P* ≤ 0.01; *** *P* ≤ 0.001

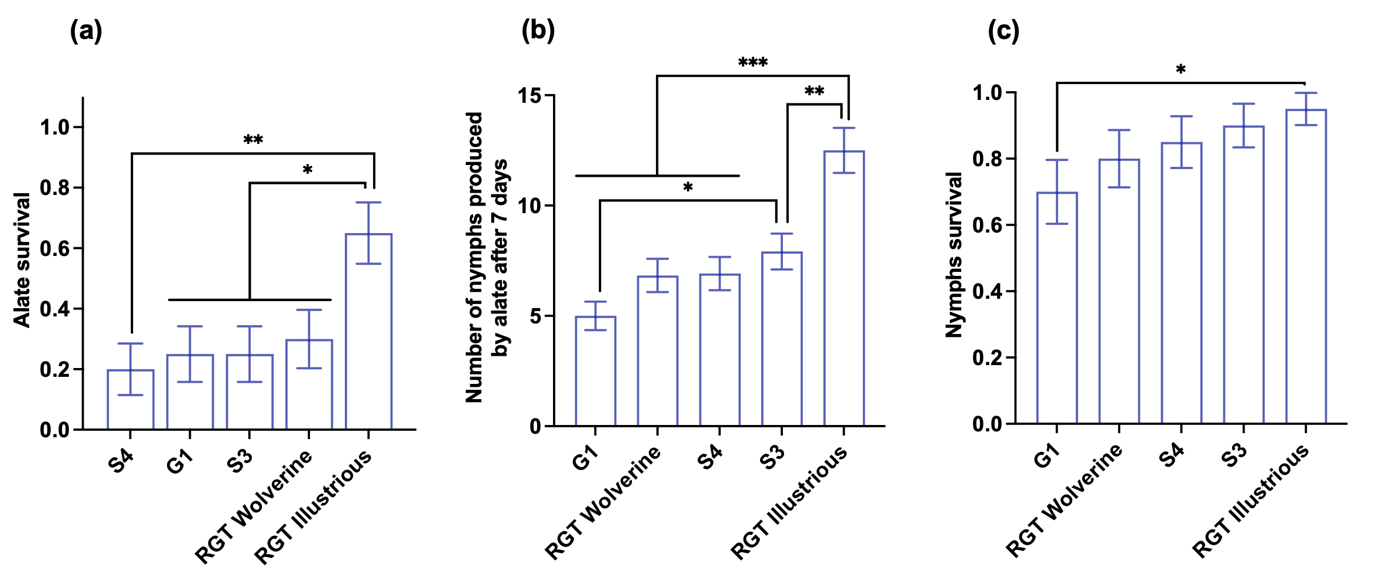

**Figure S2.** Life history traits of *R. padi* with BYDV analysed using generalised linear model (GLM) (*n* = 12). (a) Alate survival after seven days analysed with binomial regression and link function logit (*P* = 0.002). (b) Number of nymphs produced by alates after 7 days analysed with poisson distribution and link function log (*P* < 0.001). (c) Nymph survival after seven days analysed with binomial regression and link function logit (*P* = 0.027). * *P* ≤ 0.05; ** *P* ≤ 0.01; *** *P* ≤ 0.001
